# A versatile semiautomated image analysis workflow for time-lapsed camera trap image classification

**DOI:** 10.1101/2022.12.28.522027

**Authors:** Gerardo Celis, Peter Ungar, Aleksandr Sokolov, Natalia Sokolova, Hanna Böhner, Desheng Liu, John Ziker, Olivier Gilg, Ivan Fufachev, Olga Pokrovskay, Rolf Anker Ims, Valeriy Ivanov, Dorothee Ehrich

## Abstract

1. Camera trap arrays can generate thousands to millions of images that require exorbitant time and effort to classify and annotate by trained observers. Computer vision has evolved as an automated alternative to manual classification. The most popular computer vision solution is the supervised Machine Learning technique, which uses labeled images to train automated classification algorithms.
2. We propose a multi-step semi-automated workflow that consists of (1) identifying and separating bad-from good-quality images, (2) parsing good images into animals, humans, vehicles, and empty, and (3) cropping animals from images and classifying them into species for manual inspection. We trained, validated, and evaluated this approach using 548,627 images from 46 cameras in two regions of the Arctic (northeastern Norway, and Yamal Peninsula, Russia).
3. We obtained an accuracy of 0.959 for all three steps combined with the complete year test data set at Varanger and 0.922 at Yamal, reducing the number of images that required manual inspection to 7.9% of the original set from Varanger and 3.2% from Yamal.
4. Researchers can modify this multi-step process to meet their specific needs for monitoring and surveying wildlife, providing greater flexibility than current options available for image classification.

## 1 INTRODUCTION

Digital camera traps have become widely used for surveying and monitoring wildlife (Burton et al., 2015; Wearn & Glover-Kapfer, 2019). Trap arrays often generate thousands to millions of images depending on whether motion-activated or timelapse exposure is used. This requires exorbitant time and effort to classify and annotate images if done manually. Computer vision techniques have evolved as automated alternatives to manual classification, accomplishing the task at speeds greatly outpacing human capacity. The most popular computer vision solution is the supervised Deep Learning technique, which uses labeled images to train classification algorithms (Janiesch et al., 2021).

Various products to provide trained models, such as MegaDetector (Beery et al., 2019), Machine Learning for Wildlife Image Classifications (Tabak et al., 2020), and CameraTrapDetectoR (Tabak et al., 2022) have been developed to aid researchers with image classification. However, these tend to underperform manual classification when images are derived from locations where models are not trained, and several authors emphasize the need for human checking of many images (Schneider et al., 2020; Vélez et al., 2022; Fenell et al. 2022). Examples of complete workflows from image retrieval over automatic classification to manual checking and the assemblage of final metadata files with annotations have also been made available (Böhner et al., 2022; Vélez et al., 2022). While pre-trained models are often limited to the fauna of specific regions (e.g. Deepfaune; Rigoudy et al., 2022 or Conservation AI), the training of custom models may require millions of images and computer processing power inaccessible to many researchers. A more streamlined, accessible, and versatile solution that utilizes the strengths of computer vision models is needed.

While many studies use cameras triggered by a motion sensor (Böhner et al., 2022), in some cases, when there is a lot of disturbance from snowfall or when motion sensor triggering is not reliable, a time-lapse protocol is more appropriate (Hamel et al., 2013). Compared to motion sensor pictures, time-lapse camera trap datasets contain many more empty pictures. For a classification aiming at minimizing hands-on time required for manual checking, a reliable identification of empty pictures is thus particularly important. At the same time, in order to maintain data quality for animals that do not remain long at bait stations, it is important to minimize false negatives, and empty pictures should be distinguished from pictures with bad visibility in order to relate detections to observation effort.

This paper presents a solution that is specifically tailored to time-lapse protocols containing a large number of empty pictures. It provides a guided example of a multi-step workflow for semi-automated classification of images from camera traps using a personal computer with a GPU or multi-thread CPU (Windows, Macintosh, and Linux operating systems). Our approach combines training custom models that can be adapted to a new context in a flexible way with a highly performant openly available model, MegaDetector. Specifically, it consists of (1) identifying good-quality images, (2) separating empty images from images with animals, humans, or vehicles (3) cropping out animals from images and classifying them by species, and (4) manual inspection of a selection of images. Fennell et al. (2022) and Rigoudy et al. (2022) used similar approaches combining Megadetector with custom trained species identification, but contrary to their case, in our study limiting false negatives was crucial. We investigate optimal thresholds and procedures for the trade-off between false negatives and manual reviewing time.

We demonstrate the workflow by applying it to two examples from the Arctic, representing sampling at a total of six sites in the Yamal Peninsula, Russia, and northeastern Norway. Our datasets consist of time-lapse images taken at bait stations. Classification of camera trap images from the Arctic is especially challenging given ever-changing backgrounds due to seasonal extremes in snow, vegetation cover, and light conditions (day length), thus providing an excellent test case to show the potential of this approach. Furthermore, because arctic ecosystems are at present rapidly changing under the impact of climate change and increasing human activity, there is an urgent need for thorough monitoring of important species such as carnivores, and camera traps are a well-suited non-invasive method that can relatively easily be deployed in remote areas (Hamel et al., 2013).

## 2 MATERIALS AND METHODS

### 2.1 Camera trap setup and datasets

Images were collected from camera traps used in a monitoring program of the tundra carnivore scavenger guild in two low Arctic regions, the Yamal Peninsula, Russia, and northeastern Norway (Killengreen et al., 2012). In Yamal, ten cameras were deployed at one site, Erkuta (68.2° N, 69.1° E), and in Norway 36 cameras were spread across five sites (Komagdalen, Vestre Jakobselv, Stjernevann on Varanger Peninsula, and Ifjordfjellet and Gaissene in Laksefjordvidda; 70-71° N, 25-30° E; hereafter we refer to the whole study area in Norway as Varanger) (Table 1). Cameras were activated during late winter from the end of February to early April and data used in this study were collected from 2016 to 2022.

**Table 1.** The number of cameras deployed in the field for each site and the number of images used for model training, validation, and testing from Varanger, Norway (2016-2018, 2020-2021) and Yamal, Russia (2017-2021), and out-of-sample validation (Varanger 2019, and Yamal 2022).

| Region | Site | Number of Cameras | Images for models | Image for Out-of-Sample |
| --- | --- | --- | --- | --- |
| Varanger | Komagdalen | 8 | 461,407 | 75,824 |
|  | Vestre Jakobselv | 7 | 367,483 | 79,807 |
|  | Stjernevann | 5 | 284,994 | 53,555 |
|  | Ifjordfjellet | 8 | 401,039 | 58,652 |
|  | Gaissene | 8 | 426,970 | 78,620 |
| Yamal | Erkuta | 10 | 535,910 | 120,840 |

We used RECONYX® cameras (Rapid Fire, Hyperfire and Hyperfire 2, Holmen, WI, USA) placed on a permanently fixed metal pole at 30-50cm above the snow surface. Cameras were painted in white and equipped with external batteries. In Varanger, each camera station was baited with a ca 15 kg frozen block of reindeer meat and in Yamal frozen pelvis bones of reindeer with 1-2 kg remains of flesh (or muscle tissue) was mounted on a metal pole placed 2-5 m north of the camera in the direction of its optical view. Cameras were programmed to take a picture every 5 minutes (no motion sensor). After 2-3 weeks of deployment, baits were replaced if needed, and memory cards were collected and replaced until the end of the sample period.

Initially, all images were reviewed manually using the software MapView Professional (RECONYX ®) and separated into bad quality images (*Bad*) that are out of focus or obstructed (snow/ice in front of the lense or snowstorms), and good quality images where an animal could have been detected (*Good*). *Good* images were classified by animal presence/absence (*Animal*), and species and number of the animal(s), when present. A total of 2,285,351 images were generated in Varanger (2016-2021) and 656,750 images in Yamal (2017-2022; Table 1). The vast majority of images from both locations were classified as *Good* (>83%). In Varanger, *Bad* images represented 18.1% and in Yamal *Bad* images represented 8.7% (Table 2). At least one animal was detected in 6.9% of the images from Varanger and in 2% of the images in Yamal.

**Table 2.** The total number of images per classification group as assessed manually at Varanger and Yamal. Median and mean (standard deviation) percentage of images for each individual camera trap per year at Varanger, (N = 36) and years (2016 - 2021) for a total of 2,285,351 images and at Yamal (N = 10) and years (2017 - 2022) total of 656,750 images.

| Location | Class ID | N | Median | Mean (SD) |
| --- | --- | --- | --- | --- |
| Varanger | Bad | 402,409 | 14.5 | 18.1 (14.5) |
|  | Good Animal | 150,532 | 6.5 | 6.9 (3.5) |
|  | Good Empty | 1,735,410 | 78.5 | 75.7 (13.5) |
| Yamal | Bad | 55,420 | 5.4 | 8.7 (9.4) |
|  | Good Animal | 14,133 | 1.3 | 2.0 (1.7) |
|  | Good Empty | 587,197 | 93.4 | 90.1 (9.3) |

Twelve species of mammals and birds were documented in both locations, although community structure differed between the two (Table 3). In Varanger, the most common species was the raven (*Corvus corax*), appearing in 121,409 images, followed by red fox (*Vulpes vulpes*) in 14,903 images and golden eagle (*Aquila chrysaetos*) in 5,168 images. In Yamal, the most common species was the Arctic fox (*Vulpes lagopus*), appearing in 5,017 images, followed by the magpie (*Pica pica*) in 4,269 images and the mountain hare (*Lepus timidus*) in 1,677 images.

**Table 3.** The total number of appearances for each species as assessed manually at Varanger for all camera traps (N = 36) and years (2016-2021) and Yamal (N= 10; 2017-2022). A total of 148,342 images for Varanger and 14,332 for Yamal contained animals.

| Class ID | Varanger | Yamal | Included in model |
| --- | --- | --- | --- |
| Moose - *Alces alces* | 66 | 0 | Yes |
| Golden eagle - *Aquila chrysaetos* | 5,168 | 0 | Yes |
| Snowy owl - *Bubo scandiacus* | 69* | 2 | No |
| Dog – *Canis familiaris* | 0 | 19 | No |
| Raven - *Corvus corax* | 121,409 | 246 | Yes |
| Hooded crow - *Corvus cornix* | 1,011 | 38 | Yes |
| Wolverine *Gulo gulo* | 1,103 | 171 | Yes |
| White-tailed eagle - *Haliaeetus albicilla* | 1,474 | 0 | Yes |
| Human – *Homo sapiens* | 152 | 1,044 | No |
| Ptarmigan – *Lagopus* spp.** | 0 | 131 | Yes |
| Mountain hare – *Lepus timidus* | 0 | 1,677 | Yes |
| Magpie - *Pica pica* | 1 | 4,269 | Yes |
| Reindeer - *Rangifer tarandus* | 3,143 | 679 | Yes |
| Arctic fox - *Vulpes lagopus* | 2,341 | 5,017 | Yes |
| Red fox - *Vulpes vulpes* | 14,903 | 1,037 | Yes |

\* All images were from the out-of-sample data set of the year 2019, this species was thus not used to train the model.
\*\* Most ptarmigan observed in Erkuta are willow ptarmigan *Lagopus lagopus* but rock ptarmigan *Lagopus muta* occur as well, but it is difficult to identify them reliably on camera trap pictures. Both species are also present on Varanger, but they were not recorded systematically in that data set.

#### 2.1.1 Workflow test data set

Although cameras were placed at the same location every year, the background within the site varied both within and between years (e.g snow cover and exact camera positioning), creating distinct image sets (Fig. S2). This added complexity to the images allowed us to test our workflow (Fig. 1) under challenging “real-world” conditions. Therefore, we set aside one complete year of images from each site to use as test data sets that were not included in the training and validation data sets. This approach corresponds to the situation of long-term monitoring programs, where new image datasets are obtained annually and should be classified with a procedure developed based on available data from previous years (Böhner et al. 2022). We used images from all cameras in Varanger in 2019 (images per site: Komagdalen 75,726, Vestre Jakobselv 79,714, Stjernevann 53,532, Ifjordfjellet 58,504, and Gaissene 78,460) and Erkuta in 2022 (120,840 images) to test our whole workflow (Table 1).

**Figure 1.**
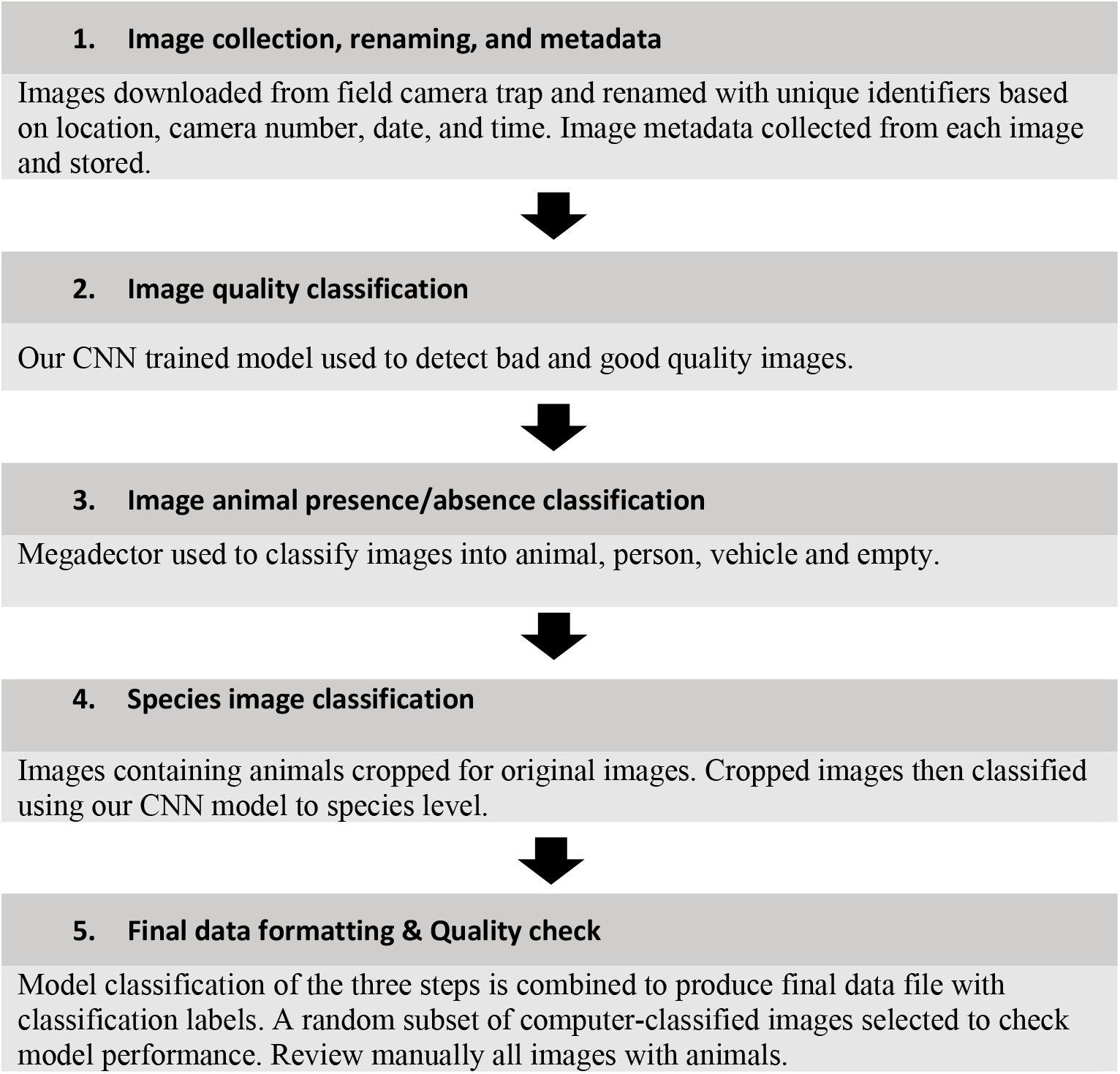
Time-lapse camera trap workflow. Data preparation and model steps are adapted from (Böhner et al., 2022).

### 2.2 Workflow

The multi-step semi-automated workflow proposed here (Fig. 1) is adapted from Böhner et al. (2022), including pre-processing of images, model training, manual quality checks, and final data formatting. Specifically, we build on the results of Rigoudy et al. (2022) and Fennell et al. (2022) who combined Megadetector with manual classification by combining custom trained models with Megadetector. The workflow consists of the following 3 steps in addition to pre-processing of images and final data formatting (Fig. 1).

#### 2.2.1 Step 1: Image quality classifications

As an initial step we were interested in creating an image classification that could parse images by image quality. Separating empty images from bad quality images is important to quantify the observation effort as number of good quality images per day for downstream modelling of detections. We developed a model that could classify images as *Bad* or *Good* quality.

Based on the manual classifications of images we randomly selected images from each site, camera, and year, such that we had a ∼15k images of *Bad* quality and ∼57k images of *Good* quality for each location on Varanger and Yamal. All of these images were then reexamined by GC and DE, and any images misclassified by the original observer were removed. We also excluded difficult images (e.g. partly blurred images, images where an animal is only visible with a tail in a corner etc.), as a good quality training data are important for model training (Böhner et al., 2022). In particular, images of animals at extreme distances (e.g., appearing as points on the horizon more than 3 m from the bait station) that could be identified by humans only because they moved in and out of frame were excluded from the model training.

The resultant data subset was randomly divided into 90% to be used for the training model, 8% for validation internally in model training, and 2% for evaluation of the trained model. We recommend this data split if user trains their own models. A total of 47,452 images were used for

Varanger and 34,399 for Yamal after manual reexamination (Table S1). These datasets were then used for all subsequent steps of the workflow. After evaluating a range of confidence thresholds for each classification class, we found that best results were obtained by using a 0.9 threshold for *Bad* quality class in the image quality model

#### 2.2.2 Step 2: Animal presence/absence classifications

For animal detection we usedMegaDetector v5.0b (https://github.com/microsoft/CameraTraps/; Beery et al., 2019), which can also detect humans and vehicles present in each image. MegaDectector was created to find and localize objects in images by creating a bounding box. In our case, MegaDetector proved superior to any other existing product at detecting animals. It further allows cropping individual animals from images using the bounding boxes, which reduces pixels to only include animals, thus facilitating determination of species identity and individual abundance.

We used asymmetric confidence thresholds or confidence probability of prediction for a class for the model classifications to reduce the number of false negatives and rather increase the number of false positives. We applied a 0.1 confidence threshold for Animal, 0.2 for humans and 0.8 for vehicles. Human and vehicle classes were combined because manual assessment treats the two classes as the same.

#### 2.2.3 Step 3: Species classification

After classifying images using MegaDetector, we used the bounding boxes of each object detection in the image to crop individual images with pixels that contain only the object (Fig. S1). Each bounding box had an associated confidence value (0-1), which is the probability of the model’s correct prediction of the class (animal, human, or vehicle). We considered only those classified by the model as animals for cropping using a confidence threshold greater than or equal to 0.1 and identified to the species level. Using the threshold of 0.1 resulted in images also included rocks, baits, and empty crops. We include all crop classes that had more than 50 images for a given class from the combined sites (Varanger and Yamal) and did not include any with humans or vehicles. We obtained 42,591 images to train the model, 3,746 for validation, and 939 to evaluate at the species level (Table S2).

We used the maximum model confidence value of all classes for a particular image to determine the class for that image.

### 2.3 Model training

We developed two detection tracks: one for image quality (2 classes), and the other to identify species (16 classes, Tables S1 & S2). Both models used the same protocol but each had a different data set for training, validation, and evaluation.

Models were trained in R (Team, 2022) using *keras* package (Allaire & Chollet, 2022) with a Tensorflow backend (Allaire & Tang, 2022). The ResNet50 architecture (He et al., 2015) was used to train the models with 55 epochs (number of times the algorithm goes through the entire training data set) and 64 batch size (number of samples to work through before updating model parameters) with a one-cycle learning rate (hyperparameter controlling model response to estimated error each time the model weights are updated) policy with a minimum of 0.000001 and a maximum of 0.001 (Smith, 2018) using separate training and validation image data set for each image collection region.

We used the *keras* image_data_generator function for real-time image augmentation, which included random assignment of the following: rotation 0-40 degrees, width and height shift range of 20%, shear range 0-0.2 radians, zoom range 0-0.2 scalar range, a horizontal flip and a fill mode with the nearest pixel.

We ran models locally on a laptop (MacBook Pro, M1 Pro 8-core central processing unit - CPU, 14-core graphics processing unit - GPU 16GB of RAM memory) using the GPU rather than CPU for data processing. GPUs are optimized for complex imaging tasks and in our case outperforms CPU by ∼7x in time required to process models.

### 2.4 Model performance

Model performance was assessed using the evalution data sets. We also determined the performance of our workflow using the out-of-sample data set representing a complete season of data from each site. Performance was measured in terms of accuracy, precision, recall, and F1 metrics as defined in Table 4 using *caret* R package (Kuhn, 2022). For the out of sample dataset, we also compared the number of days with detection between the worfkflow results and the manual scoring, in addition to the picture-by-picture performance evaluation. Indeed, daily counts or detections are often used in downstream analyses of camera trap data (Hamel et al., 2013; Rød-Eriksen et al., 2022).

**Table 4.** Model performance metrics based on “caret” R package. True positives (TP), true negatives (TN), False positives (FP), False negatives (FN)

| Metric | Equation | Definition |
| --- | --- | --- |
| Accuracy | $\frac{TP + TN}{TP + FP + TN + FN}$ | Proportion of correct prediction in the whole data set |
| Precision | $\frac{TP}{TP + FP}$ | The probability of the specific class is correctly classified as present given the model classified it as present. |
| Recall | $\frac{TP}{TP + FN}$ | The probability of the specific class is correctly classified by the model given that the class is truly present. |
| F1 | $\frac{2 * precision * recall}{precision + recall}$ | Weighted average of precision and recall. Represents the balance of correct prediction for all considered classification. |

## 3 RESULTS

### 3.1 Trained Model Performances

#### 3.1.1 Image Quality Models

Image quality model performance for Varanger and Yamal had accuracies of 0.995 and 0.998, respectively (Table S3). Both models had high precision (Yamal = 0.987, Varanger = 0.983) and high probability that the class is correctly retrieved by the model given that the class is truly present (Yamal = 0.996, Varanger = 1.000) for the bad quality class (Table S3, Fig. S3).

#### 3.1.2 Species Model

Species model performance had an accuracy of 0.970 (Table S3). All model class predictions evinced high precision > 0.929 with the lowest being magpie and the Arctic fox (Table S3). Recall was also high for most classes except for white-tailed eagle (0.636) and wolverine (0.889). The low recall is because 36.4% of images with a white-tailed eagle were incorrectly classified as golden eagle. The two species are indeed very similar in appearance, especially when not all animal features are visible or lighting is low. For wolverines, 11.1% were incorrectly classified as arctic fox (Fig. S4). These images were taken in the dark and mostly showed only a little distinct part of the animal, such as the back. In such cases, manual scoring often uses consecutive images to identify the species correctly.

### 3.2 Workflow performance

#### 3.2.1 Step 1 | Image Quality Model

Image quality models for out-of-sample data sets had high accuracy for both Varanger (0.970) and Yamal (0.959), though Varanger had higher precision (0.904) and recall (0.925) for bad-quality images than Yamal (0.746 and 0.794, respectively) (Table 5). There is a balance between the number of false negatives for both bad and good classes (Fig. S5), which indicates that there are some images that are borderline between good and bad for which both manual observation and the model can have different classifications. For example, images with a partial lens obstruction (e.g., snow), which the model would classify as good but manually as bad. A total of 47,433 (13.7%) bad-quality images were identified in Varanger and 11,194 (9.3%) in Yamal, both percentages within the range of manual bad-quality classifications (Table 2).

**Table 5.** Model performance metrics for classes predicted by the image quality model with out-of-sample data set at Varanger and Yamal.

| Location | Id | Precision | Recall | F1 |
| --- | --- | --- | --- | --- |
| Varanger | Bad | 0.904 | 0.925 | 0.914 |
|  | Good | 0.988 | 0.985 | 0.986 |
| Yamal | Bad | 0.746 | 0.794 | 0.769 |
|  | Good | 0.972 | 0.963 | 0.967 |

#### 3.2.2 Step 2 | Megadector model

##### Varanger data set

After eliminating the bad-quality images, the Megadetector model had an accuracy of 0.909, where animal prediction class had a low precision of 0.470, but a high recall of 0.980 (Table 6 & Fig. S6). The empty class had a precision of 0.998 and 0.903 recall. Humans had low precision and recall < 0.167, where images containing humans were of people kite-surfing at a distance not recognizable by the model. A total of 50,667 (14.6%) images were classified as having animals present, but approximately half of those were empty (Fig. S6).

**Table 6.** Model performance metrics for classes predicted by the Megadetector model with out-of-sample data set at Varanger and Yamal.

| Location | Class id | Precision | Recall | F1 |
| --- | --- | --- | --- | --- |
| Varanger | Animal | 0.470 | 0.980 | 0.636 |
|  | Empty | 0.998 | 0.902 | 0.948 |
|  | Human | 0.057 | 0.167 | 0.085 |
| Yamal | Animal | 0.135 | 0.784 | 0.230 |
|  | Empty | 0.993 | 0.891 | 0.939 |
|  | Human | 0.106 | 0.272 | 0.153 |

Excluding the false positive animal images, the total number of days in which an animal was present in an image are similar to that of manual classification with only one camera underestimating reindeer and two cameras underestimating red foxes (Fig. S7).

##### Yamal data set

The Megadetector model with Yamal data had an accuracy of 0.887; animal class had a low precision of 0.138, but a high recall of 0.784 (Table 6, Fig. S6). The empty class had a precision of 0.993 and recall of 0.892 and the human class had a precision of 0.106 and recall of 0.272.

The low precision and recall of the human class are due to two particular cases (609 images); one of the cameras showed a bait that Megadetector identified incorrectly as a human and the other camera was facing a utility cabin that Megadetector identified as a vehicle. A total of 12,440 (10.2%) of images were incorrectly classified as having animals present, but 10,719 of those were empty (Fig. S6).

Excluding the false positive animal images, the total number of days in which an animal was present in an image are mostly similar to manual classification except for magpies, mountain hares, and Arctic foxes (Fig. S8).

#### 3.2.3 Step | 3 Species model

##### Varanger data set

A total of 66,610 image crops were created from the Megadector model for animal classified images after removing bad-quality images. Forty-nine images were of new classes that were not included in the model (human, snowy owl, black-backed gull – *Larus marinus*). After excluding these, the accuracy for the species model in Varanger was 0.88. The model was very precise at classifying cropped images as Bait and Empty (>0.92; Table 7), with only one animal image misclassified as empty (Fig. 10). The model was less precise in classifying rocks (0.83), but 2,441 of the 2,652 were either empty or bait (Fig. S9).

**Table 7.**
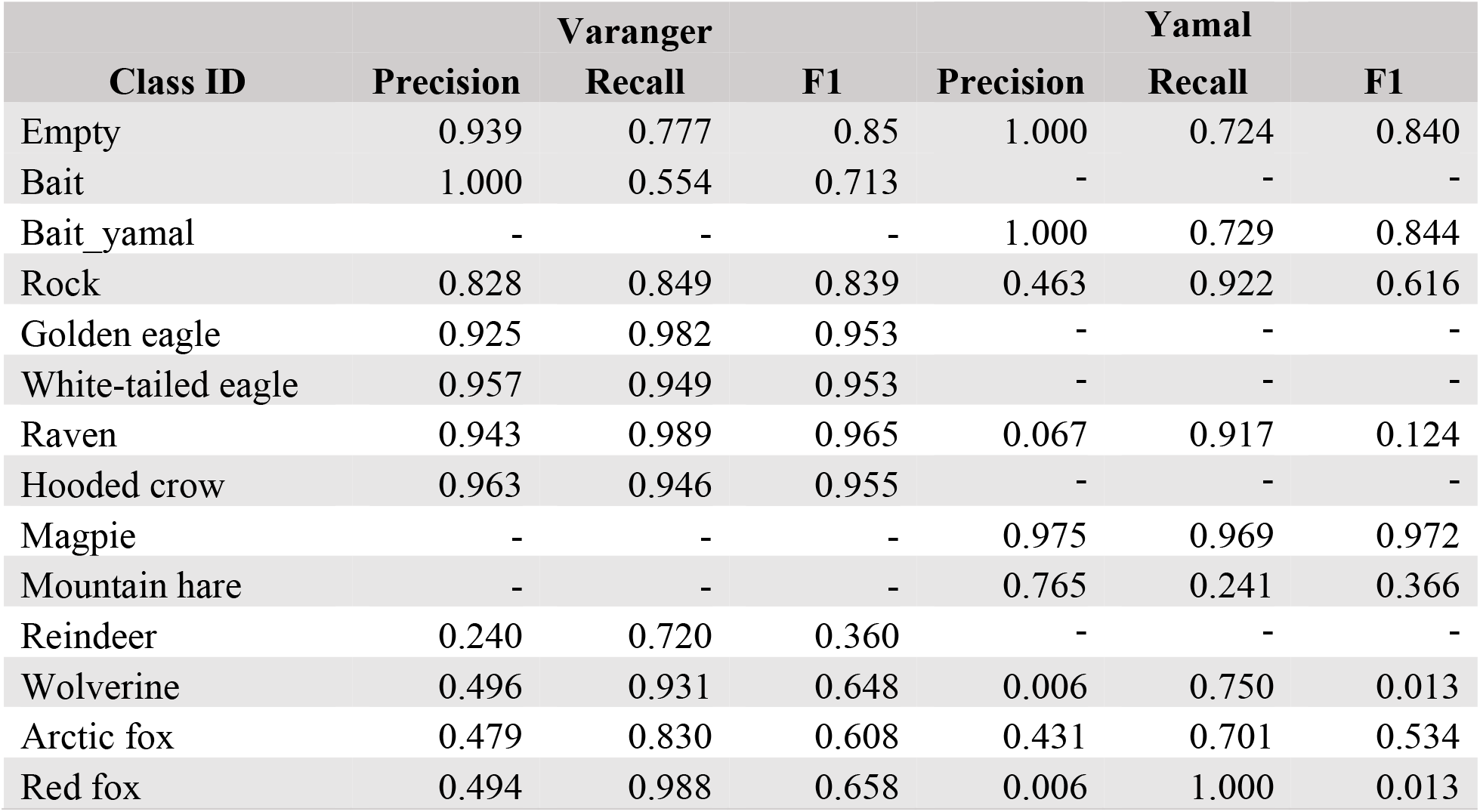
Model performance metrics for classes predicted by the species model with Varanger and Yamal workflow test data set. To estimate model performance metrics, classes that were exclusively in the manual assessment or model output were not included. See Table 3 for species names.

| Class ID | Varanger |  |  | Yamal |  |  |
| --- | --- | --- | --- | --- | --- | --- |
|  | Precision | Recall | F1 | Precision | Recall | F1 |
| Empty | 0.939 | 0.777 | 0.85 | 1.000 | 0.724 | 0.840 |
| Bait | 1.000 | 0.554 | 0.713 | - | - | - |
| Bait_yamal | - | - | - | 1.000 | 0.729 | 0.844 |
| Rock | 0.828 | 0.849 | 0.839 | 0.463 | 0.922 | 0.616 |
| Golden eagle | 0.925 | 0.982 | 0.953 | - | - | - |
| White-tailed eagle | 0.957 | 0.949 | 0.953 | - | - | - |
| Raven | 0.943 | 0.989 | 0.965 | 0.067 | 0.917 | 0.124 |
| Hooded crow | 0.963 | 0.946 | 0.955 | - | - | - |
| Magpie | - | - | - | 0.975 | 0.969 | 0.972 |
| Mountain hare | - | - | - | 0.765 | 0.241 | 0.366 |
| Reindeer | 0.240 | 0.720 | 0.360 | - | - | - |
| Wolverine | 0.496 | 0.931 | 0.648 | 0.006 | 0.750 | 0.013 |
| Arctic fox | 0.479 | 0.830 | 0.608 | 0.431 | 0.701 | 0.534 |
| Red fox | 0.494 | 0.988 | 0.658 | 0.006 | 1.000 | 0.013 |

##### Yamal data set

A total of 13,754 image crops were created from the Megadetector model classified as containing animals, after removing bad-quality images. Five images were not included in the trained model (humans, snowy owl, and ptarmigan) and 51 were erroneously classified as reindeer, which did not exist in the data set. After excluding these, the overall accuracy for the species model in Yamal was 0.745. The model was very precise at classifying image crops that contained a bait or that were empty (Bait_yamal and Empty, 0.998; Table 7) with only one animal (wolverine) image misclassified (Fig. S9). The model was less precise in classifying rocks (0.462), but 144 of the 165 false positives were either empty or baits.

#### 3.2.4 All steps Combined

After combining all model predictions (image quality, Megadetector, and species) we obtain an overall classification accuracy of 0.959 for Varanger and 0.922 for Yamal. Including the species classification model reduced animal false positives created by Megadetector from 25,941 to 4,112 for the Varanger data set and from 10,719 to 2,942 for the Yamal data set (Figs 2 & S6). This equates to animal detection of 4,582 images or 3.8% of the total images from camera traps in Yamal, and 27,379 or 7.9% in Varanger, which are higher than the mean manual animal detection rates of 2.0% in Yamal and 6.9% in Varanger (Table 2).

**Figure 2.**
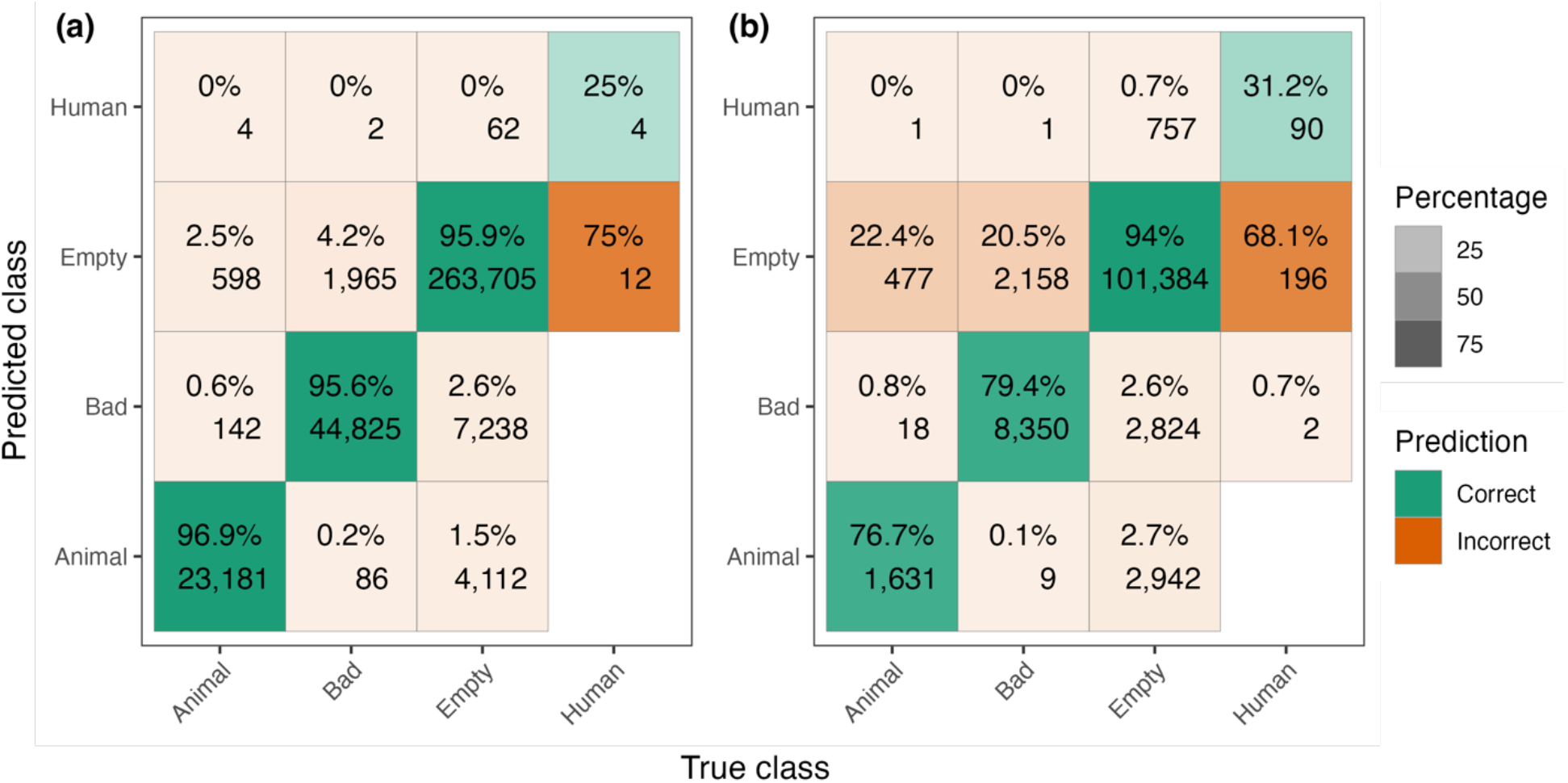
All models combined confusion matrix with (a) Varanger and (b) Yamal out-of-sample data set. Species model results were aggregated in the “Animal” class. The percentage and number of images with correct (diagonal, green) and incorrectly predicted classes (off-diagonal).

## 4 DISCUSSION/CONCLUSION

Our models correctly classified 97% of images into good/bad, 90% of images into animal/no animal, and 98% of images into the correct species when compared with a manually derived classification. We obtained an accuracy of 0.959 for Varanger and 0.922 for Yamal with low false-negative rates of 0.2% and 0.4% respectively. Assuming that all pictures where animals were detected would be reviewed manually, this would reduce the number of images that require manual inspection to 7.9% in Varanger and 3.2% in Yamal by eliminating bad-quality and empty images. Implementing this procedure could therefore save a great deal of time and effort associated with manual inspection/classification of imagery.

The proposed semi-automatic workflow for classifying camera trap images is a robust method for identifying good-quality images, identifying animals in images, and classifying them to species in both Yamal and Varanger. Our workflow detected bad-quality images and those with animals within the ranges of those detected by manual classification (9-14% and 3-8%, respectively). The species classification step reduced the number of false positive animal detections generated by Megadetector, which was initially intended to be high as a result of having a low confidence threshold (0.1 in our case) aimed at minimizing false negatives. This strategy maximizes recall (increasing the probability of detecting an animal when it is present), but it comes at the expense of lower precision (Vélez et al., 2022). Although the species model reduced the number of false positives, we recommend that users manually review images with animals because the model was not sufficiently accurate to rely solely on computer classification for species classification (hence, we call this a “semi-automated” image analysis procedure).

This is usually the case for species classification models where variable model performance is observed for different species in the model (Böhner et al., 2022; Whytock et al., 2021). As more data become available, the model can be improved by retraining it with new images (Böhner et al., 2022).

Another aspect of improving model performance is the positioning of the cameras in the field. We found that the positioning of cameras can influence the detection of false positives. We recommend that cameras not be placed to include features (e.g. rocks or building structures) in the field of view that could be misclassified as animals.

We provide code for this multi-step process in the supplemental files so that researchers can use and modify it to meet their needs for monitoring and surveying wildlife. Because our workflow is subdivided into several steps, it is flexible and can be adapted to various situations. The initial step could, for instance be modified to include a classification into pictures with and without bait in addition to quality, or with and without snow depending on the aims of the study.

## AUTHOR CONTRIBUTIONS

GC, DE AS, NS, HB, RAI, IF, and OP conceived the ideas and designed methodology; DE, AS, NS, IF, and OP collected the data; GC and HB were involved in software, validation, formal analysis; RAI, DL, JX, PSU, and VI project administration and funding acquisition; GC led the writing of the manuscript and visualization. All authors contributed critically to the drafts and gave final approval for publication.

## DATA AVAILABILITY STATEMENT

The original pictures from Varanger can be obtained from the authors upon request. The dataset from Varanger will be available from the database of the Climate-ecological Observatory for Arctic Tundra (https://data.coat.no/). The code and detailed instructions for the workflow are provided on Github (https://github.com/gerlis22/CameraTrap)

## ACKNOWLEDGMENTS

This research was made possible thanks to funding from Navigating the New Arctic – National Science Foundation (Award 2126796). The collection of the Norwegian data set was financed by the Norwegian Environmental Agency. Jan Erik Knutsen, Berit Gaski and the State Nature Surveillance in Vadsø and Tana carried out the field work in Norway. AS, NS, IF and OP were supported by the Russian Ministry of Science and Higher Education program “Terrestrial ecosystems of northwestern Siberia: assessment of the modern transformation of the communities” No. 122021000089-9. We thank the families Laptander, Serotetto and Vanuy’to who helped us in all years in challenging early spring conditions in Yamal, and Vyacheslav Osokin for unvaluable contribution to field work. Numerous assistants contributed to the manual scoring of the images, notably Torunn Moe, Stijn Hofhuis, Dag A. H. Olsen and Fanny Berthelot.

