## Supplemental table and figures for "A versatile semiautomated image analysis workflow for time-lapsed camera trap image classification"

**Supporting Information**

Table S1. Number of images used for training, validation, and evaluation for the first custom model of the workflow identifying bad quality images. For Varanger, images from the years 2016, 2017, 2018, 2020, and 2021 were used, and for Yamal we used images from the years 2017 - 2021.

| Model | Class ID | Train | Validate | Evaluation |
|---|---|---|---|---|
| Varanger | Bad | 10,678 (25.0%) | 1,050 (27.5%) | 239 (24.9%) |
| | Good | 31,993 (75.0%) | 2,770 (72.5%) | 722 (75.1%) |
| | **Total** | **42,671** | **3,820** | **961** |
| Yamal | Bad | 7,066 (22.2%) | 318 (15.6%) | 78 (15.3%) |
| | Good | 24,791 (77.8%) | 1,714 (84.3%) | 432 (84.7%) |
| | **Total** | **31,857** | **2,032** | **510** |

Table S2. Number of images used for training, validation, and evaluation for the second custom model of the workflow identifying species. Images from years 2017, 2018, 2019, 2020, and 2021 from Varanger and Yamal.

| Class ID | Train | Validate | Evaluation |
|---|---|---|---|
| Empty | 2,992 (7.0%) | 270 (7.2%) | 71 (7.6%) |
| Bait | 885 (2.1%) | 71 (1.9%) | 19 (2.0%) |
| Bait_yamal | 1,325 (3.1%) | 102 (2.7%) | 29 (3.1%) |
| Rock | 3,137 (7.4%) | 278 (7.4%) | 64 (6.8%) |
| Golden eagle - *Aquila chrysaetos* | 4,157 (9.8%) | 359 (9.6%) | 94 (10.0%) |
| White-tailed eagle - *Haliaeetus albicilla* | 590 (1.4%) | 31 (0.8%) | 11 (1.2%) |
| Raven - *Corvus corax* | 12,070 (28.3%) | 1,097(29.3%) | 247 (26.3%) |
| Hooded crow - *Corvus cornix* | 464 (1.1%) | 37 (1.0%) | 8 (0.9%) |
| Magpie - *Pica pica* | 2,323 (5.5%) | 212 (5.7%) | 39 (4.2%) |
| Ptarmigan – *Lagopus spp.* | 18 (0.04%) | 1 (0.03%) | 1 (0.1%) |
| Mountain hare – *Lepus timidus* | 61 (0.1%) | 5 (0.1%) | 3 (0.3%) |
| Moose - *Alces alces* | 24 (0.06%) | 1 (0.03%) | 1 (0.1%) |
| Reindeer - *Rangifer tarandus* | 840 (2.0%) | 59 (1.5%) | 15 (1.6%) |
| Wolverine - *Gulo gulo* | 754 (1.8%) | 74 (2.0%) | 18 (1.9%) |
| Red fox - *Vulpes vulpes* | 8,893 (20.9%) | 768 (20.5%) | 225 (24.0%) |
| Arctic fox - *Vulpes lagopus* | 4,058 (9.5%) | 381 (10.2%) | 94 (10.0%) |
| **Total** | **42,591** | **3,746** | **939** |

Table S3. Image Quality and Species Model accuracies and performance metrics for classes predicted by the image quality model with test data set from Varanger and Yamal. N is data set sample size. See table 3 for species names.

| Model | N | Accuracy | Class | Precision | Recall | F1 |
|---|---|---|---|---|---|---|
| Image Quality - Varanger | 957 | 0.995 | Bad | 0.983 | 0.996 | 0.989 |
| | | | Good | 0.999 | 0.994 | 0.996 |
| Image Quality - Yamal | 508 | 0.998 | Bad | 0.987 | 1.000 | 0.993 |
| | | | Good | 1.000 | 0.998 | 0.999 |
| Species Model | 939 | 0.970 | Empty | 0.957 | 0.944 | 0.950 |
| | | | Bait | 1.000 | 0.895 | 0.944 |
| | | | Bait_yamal | 1.000 | 0.931 | 0.964 |
| | | | Rock | 0.969 | 0.984 | 0.977 |
| | | | Golden eagle | 0.959 | 1.000 | 0.979 |
| | | | White-tailed eagle | 1.000 | 0.636 | 0.778 |
| | | | Raven | 1.000 | 0.992 | 0.996 |
| | | | Hooded crow | 1.000 | 1.000 | 1.000 |
| | | | Magpie | 0.929 | 1.000 | 0.963 |
| | | | Ptarmigan | 1.000 | 1.000 | 1.000 |
| | | | Mountain hare | 1.000 | 1.000 | 1.000 |
| | | | Moose | 1.000 | 1.000 | 1.000 |
| | | | Reindeer | 1.000 | 1.000 | 1.000 |
| | | | Wolverine | 1.000 | 0.889 | 0.941 |
| | | | Red fox | 0.957 | 0.978 | 0.967 |
| | | | Arctic fox | 0.936 | 0.936 | 0.936 |

Table S4. Performance metrics for classes predicted by all models combined with Varanger and Yamal workflow test data set.

| Location | Class id | Precision | Recall | F1 |
|---|---|---|---|---|
| Varanger | Animal | 0.847 | 0.969 | 0.904 |
| | Bad | 0.859 | 0.956 | 0.905 |
| | Empty | 0.990 | 0.959 | 0.974 |
| | Human | 0.056 | 0.250 | 0.091 |
| Yamal | Animal | 0.356 | 0.767 | 0.486 |
| | Bad | 0.750 | 0.794 | 0.769 |
| | Empty | 0.973 | 0.940 | 0.956 |
| | Human | 0.106 | 0.312 | 0.158 |

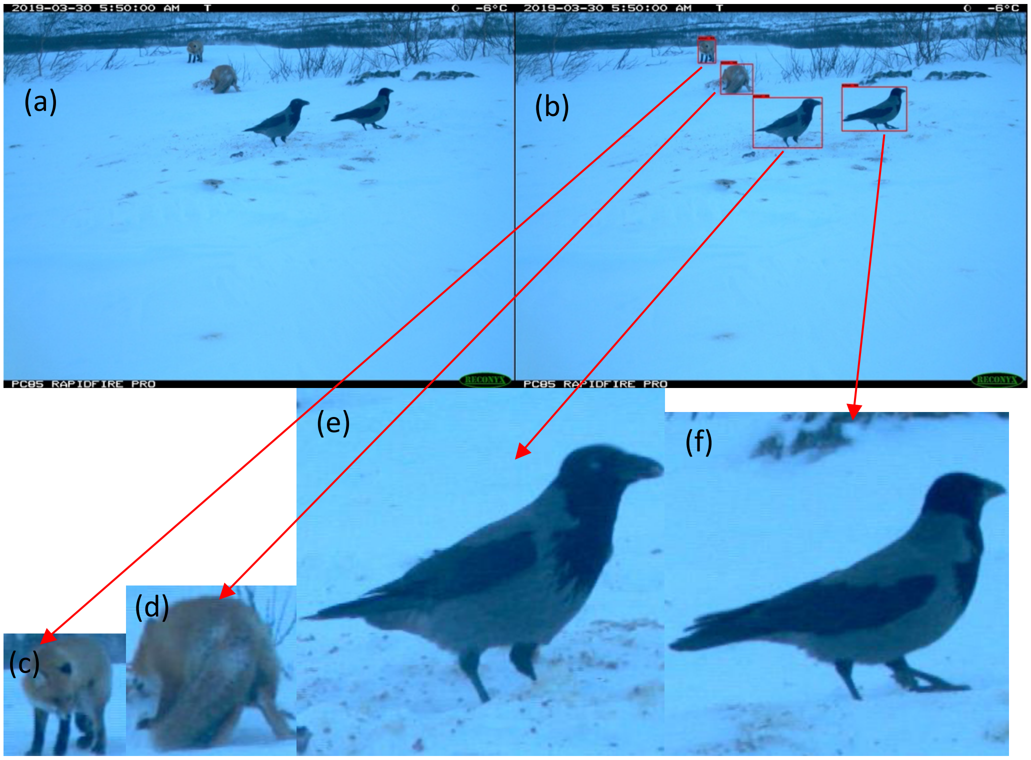


Figure S1. Camera trap image with animals present (a) and MegaDetector animal classification and bounding boxes for each detection within the image (b). Individually cropped images of animals from bounding boxes (c-f).

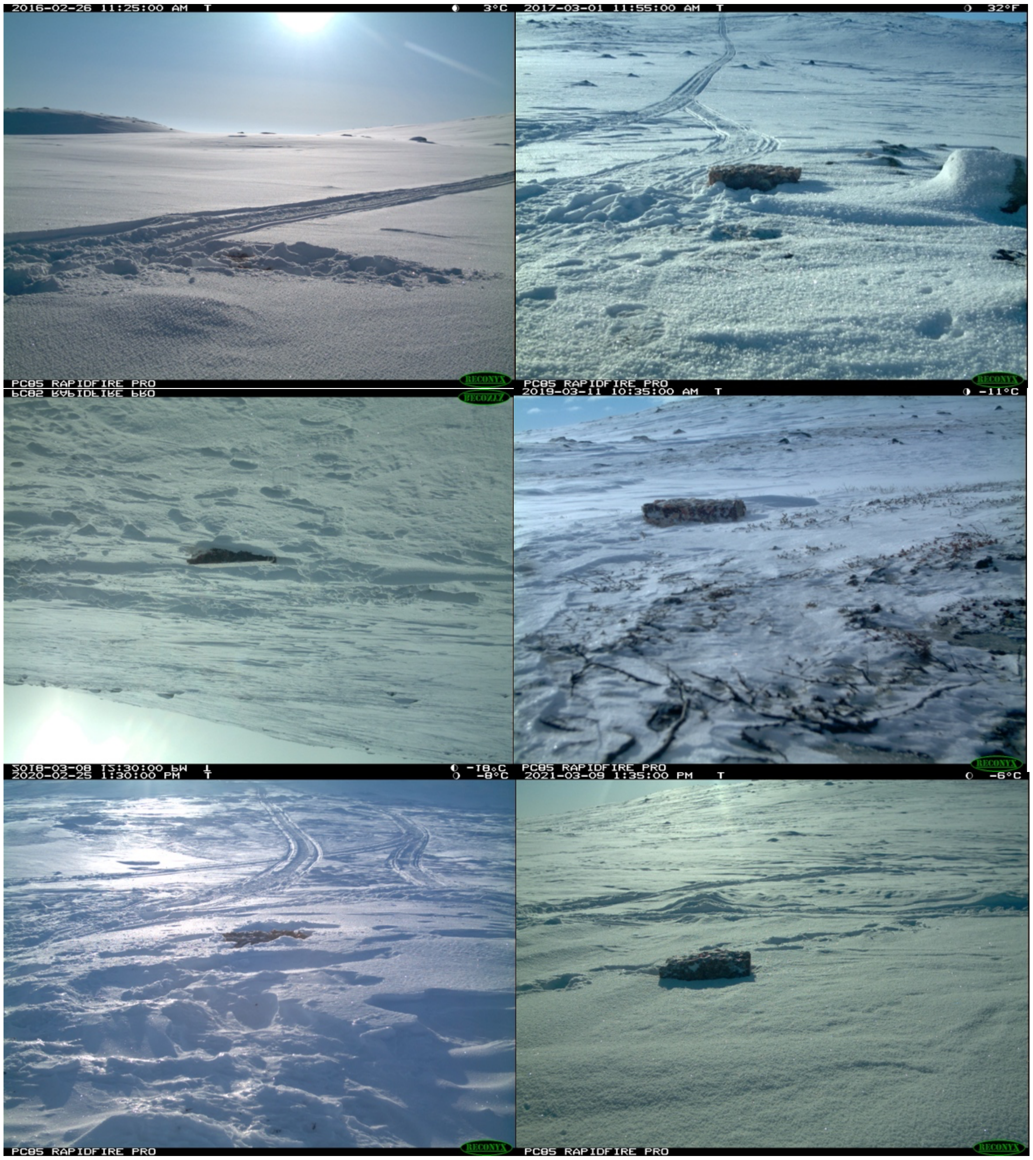


Figure S2. Varanger Camera at Varanger, Gaissene (g1) location for each year (2016-2021).

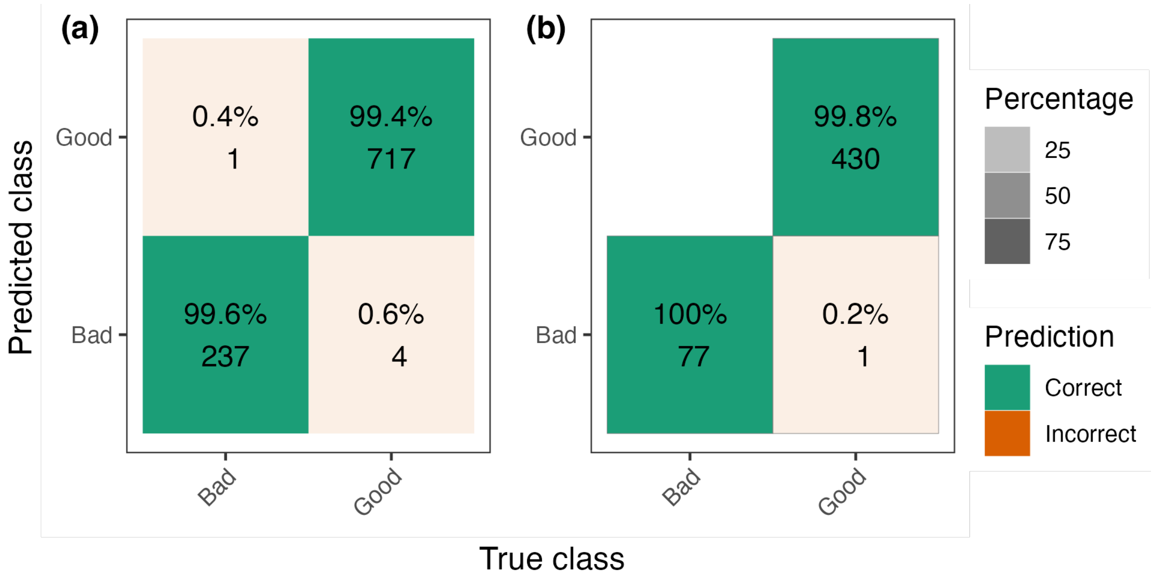


Figure S3. Image quality confusion matrix for the model evaluation data sets at (a) Varanger and (b) Yamal. The percentage and number of images with correct (diagonal, green) and incorrectly predicted classes (off-diagonal).

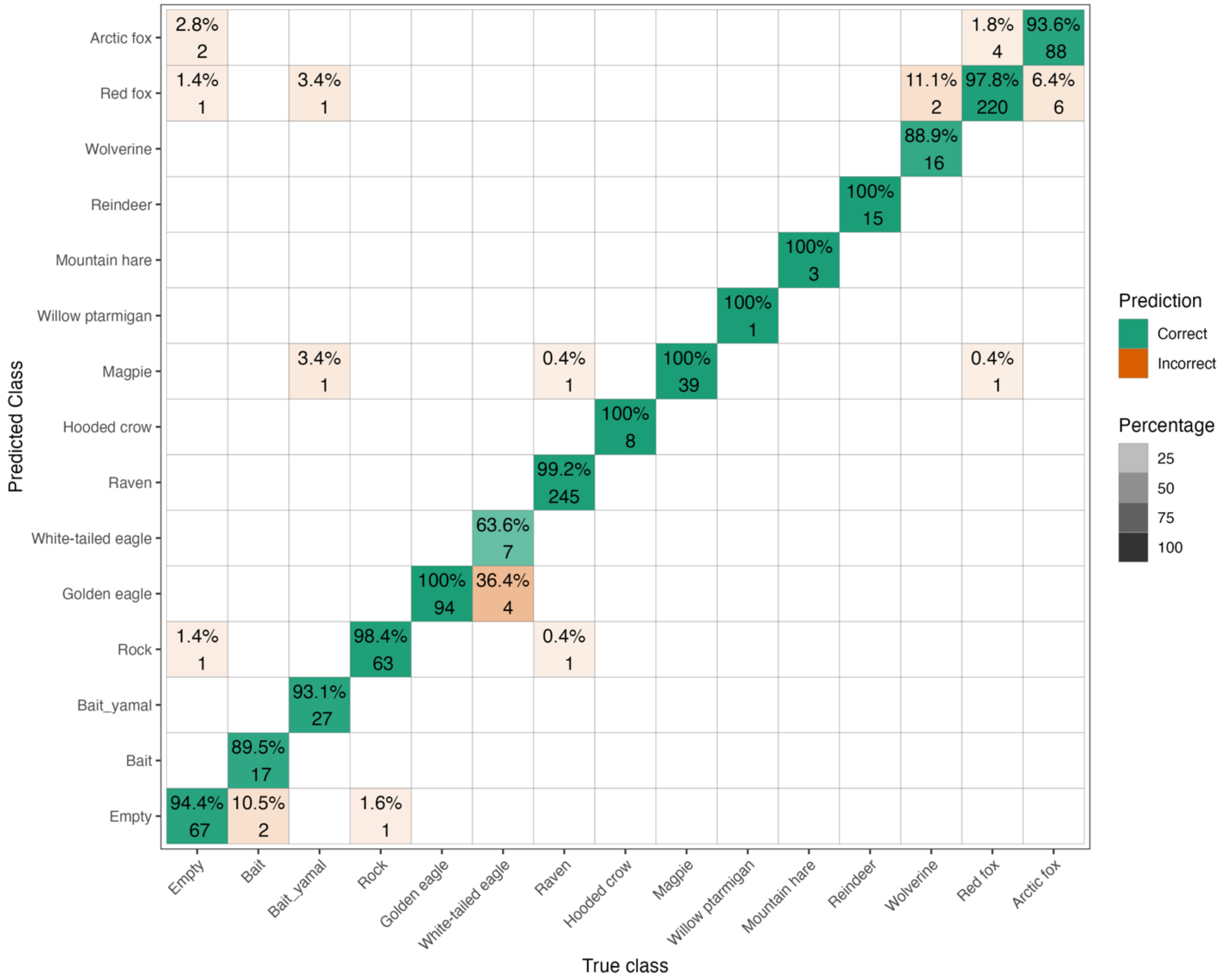


Figure S4. Species model confusion matrix for the model evaluation data set. The percentage and number of images with correct or recall (diagonal, green) and incorrectly predicted classes (off-diagonal). See table 3 for species names.

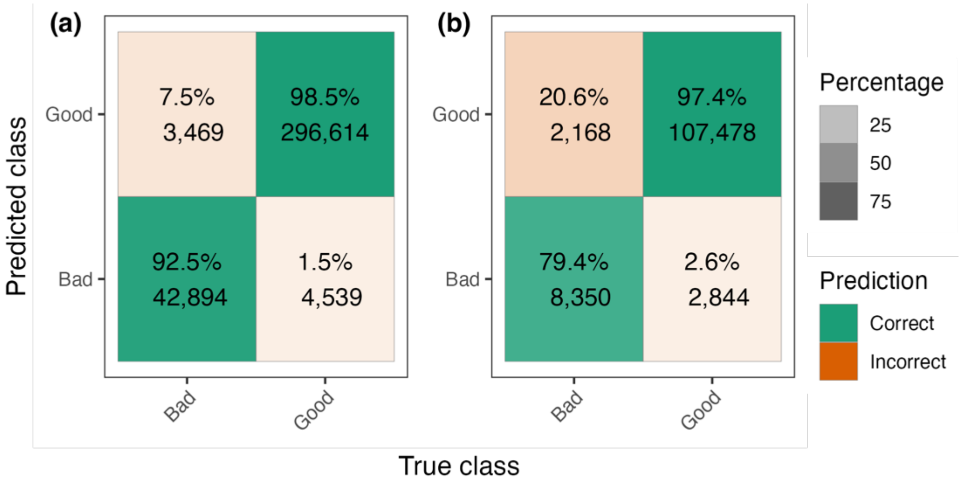


Figure S5. Image quality confusion matrix for the workflow test data set (a) Varanger and (b) Yamal. The percentage and number of images with the correct prediction (diagonal, green) and incorrectly predicted classes (off-diagonal).

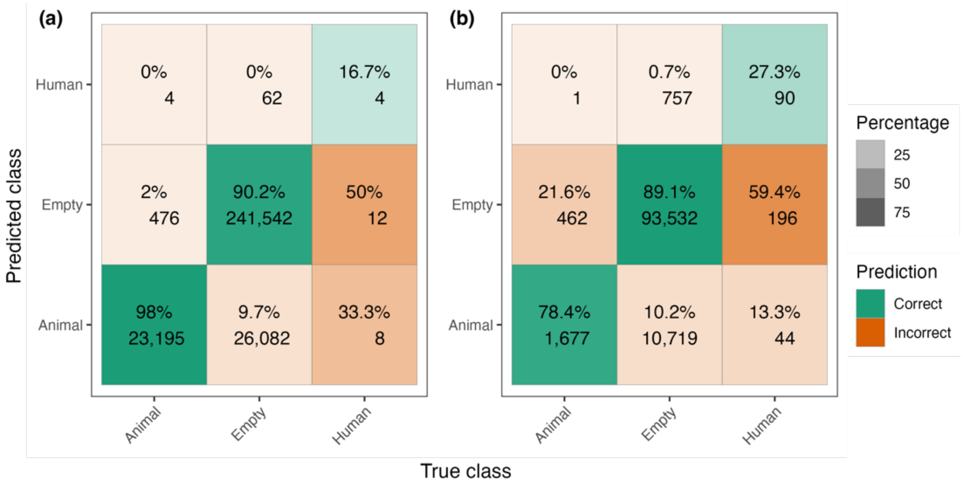


Figure S6. Megadetector model Confusion Matrix for the workflow test data set (a) Varanger and (b) Yamal. The percentage and number of images with the correct prediction (diagonal, green) and incorrectly predicted classes (off-diagonal).

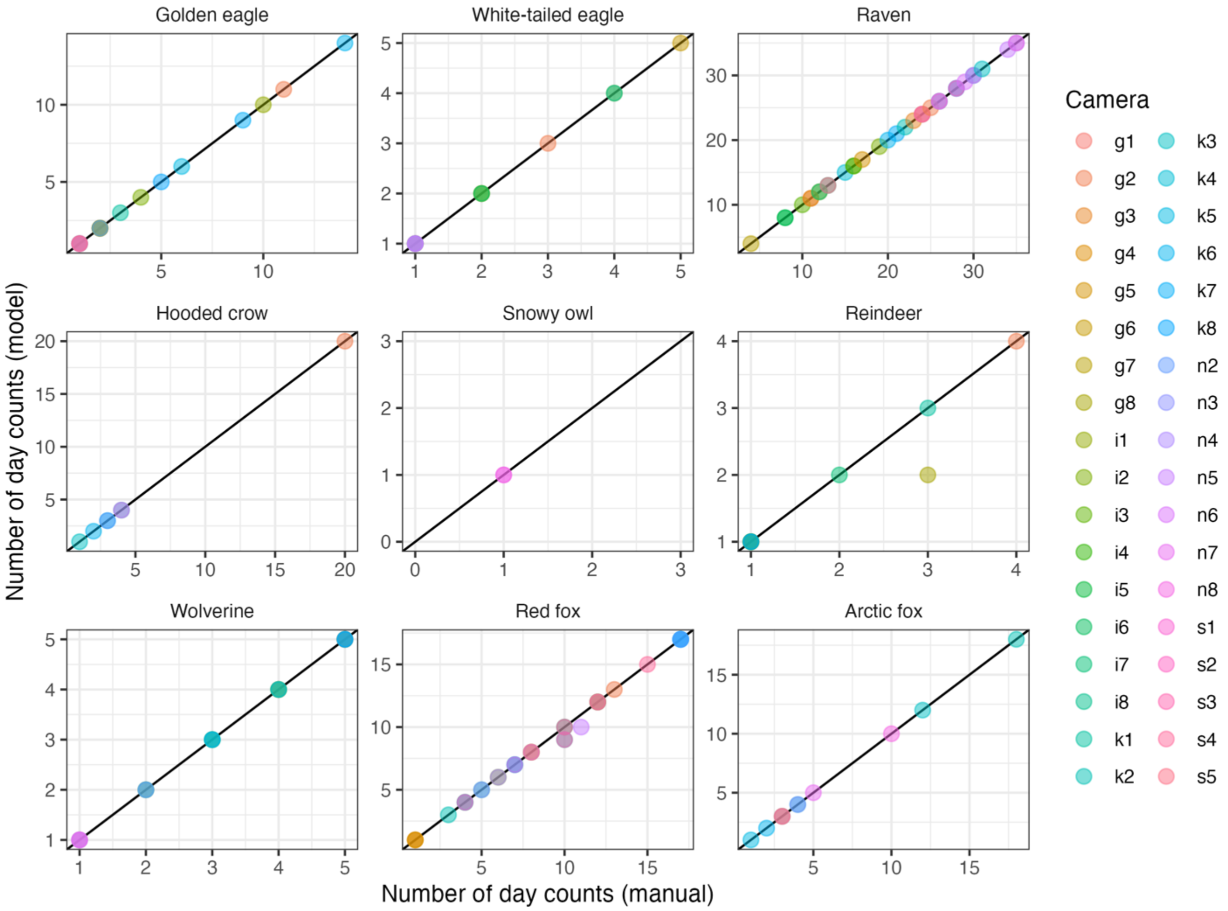


Figure S7. Number of day counts for images containing an animal with Varanger workflow test data set excluding all false positives. Black diagonal line is the one-to-one line. Each panel represents one species. See Table 4 for species acronyms.

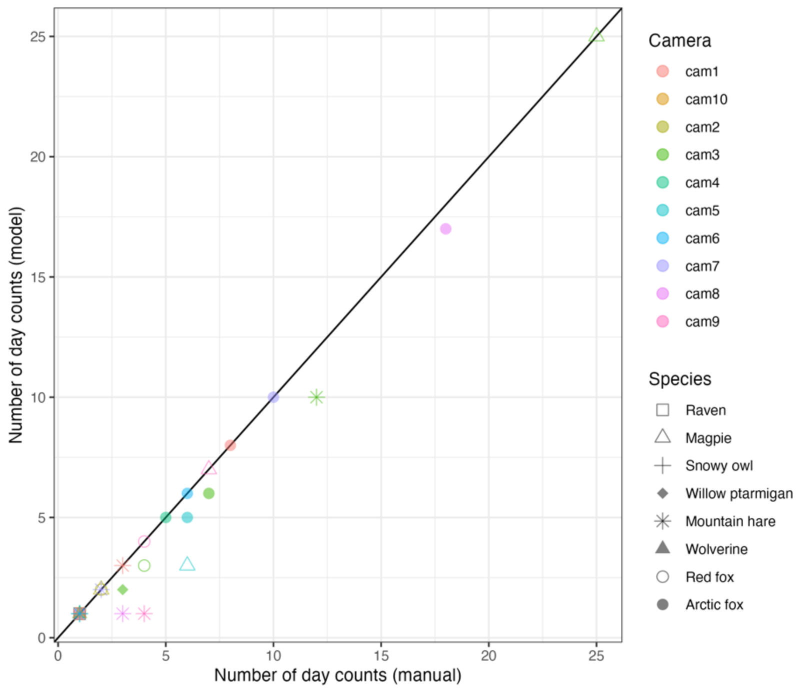


Figures S8. Number of day-counts for images containing an animal in the Yamal workflow test data set set excluding all false positives. Black diagonal line is the one-to-one line. Each shape represents a specific species. See Table 3 for species names.

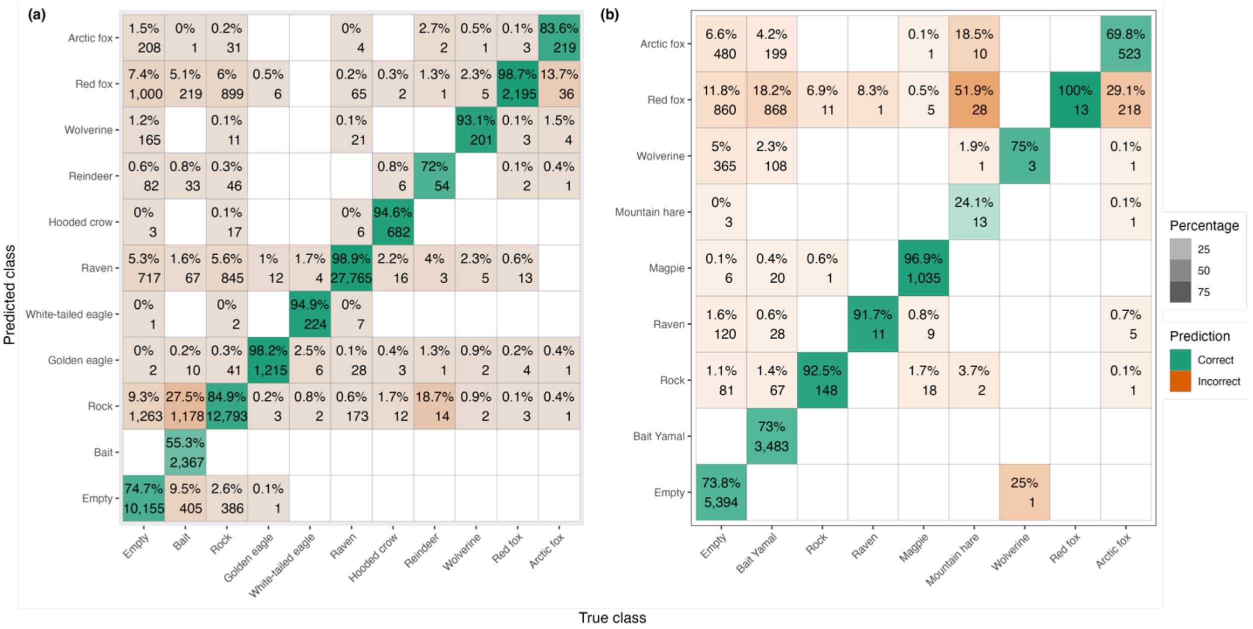


Figure S9. Species model confusion matrix for the workflow test data set (a) Varagner and (b) Yamal. The percentage and number of images with correct (diagonal, green) and incorrectly predicted classes (off-diagonal). See Table 3 for species names.
